# Optimal planning of eye movements

**DOI:** 10.1101/240010

**Authors:** Hoppe David, Constantin A. Rothkopf

## Abstract

The capability of directing gaze to relevant parts in the environment is crucial for our survival. Computational models based on ideal-observer theory have provided quantitative accounts of human gaze selection in a range of visual search tasks. According to these models, gaze is directed to the position in a visual scene, at which uncertainty about task relevant properties will be reduced maximally with the next look. However, in tasks going beyond a single action, delayed rewards can play a crucial role thereby necessitating planning. Here we investigate whether humans are capable of planning more than the next single eye movement. We found evidence that our subjects’ behavior was better explained by an ideal planner compared to the ideal observer. In particular, the location of the first fixation differed depending on the stimulus and the time available for the search. Overall, our results are the first evidence that our visual system is capable of planning.

---

Actively deciding where to direct our eyes is an essential ability in fundamental tasks, which rely on acquiring visual information for survival such as gathering food, avoiding predators, making tools, and social interaction. As we can only perceive a small proportion of our surroundings at any moment in time due to the spatial distribution of our retinal receptor cells^1^, we are constantly forced to bring task relevant parts of the visual scene into focus using eye movements^2^. Thus, vision is a sequential process of active decisions. These decisions have been characterized in terms of optimizing performance in the ongoing task^3–7^, maximizing knowledge about the environment^8–10^, or targeting gaze towards locations that are most salient^11^.

To understand the requirements of perceptual tasks, ideal-observer analysis^12,13^ has been very successful based on the idea that visual perception is inference of latent causes based on sensory signals^14,15^. In this framework, the goal of the visual system is to use sensory data *D* to infer unknown properties of the state *s* of the environment. For example, *s* could be indicating whether there is a predator hiding behind a bush, and by directing gaze to the bush visual data *D* about the latent variable describing the true state *s* of the environment is obtained. This information can be incorporated into what is known about *s* using Bayes’ theorem *P*(*s*|*D*) = *P*(*D*|*s*)*P*(*s*)/*P*(*D*). Hence, the ideal observer combines prior knowledge *P*(*s*) and sensory information *P*(*D*|*s*) to form an updated posterior belief about environmental states relevant to the specific task. The ideal-observer paradigm has been used successfully to understand how humans choose locations for the next saccade. Specifically, human eye movements use the current posterior and target the location where they expect uncertainty about task relevant variables to be reduced most after having acquired new data from that location in situations such as visual search^3^, face recognition^5^, and temporal event detection ^6^.

A limitation of ideal-observer theory is that performing sensory inference by itself does not prescribe an action, i.e. information about *s* in the end needs to be used to decide for an appropriate action, e.g. whether to flee. The costs and benefits for the potential outcomes of the action can be very different, e.g., not to flee if a predator is present is more costly than an unnecessary flight. Bayesian decision theory provides such an answer by using the costs and benefits of different outcomes with the respective uncertainties of the associated outcomes. Hence, different potential outcomes of *s* are weighted with a utility function *U*(*a, s*) to determine the action with highest expected utility: *a* = arg max*_a_ ∫_s_ U*(*a,s*)*P*(*s*|*D*)*ds*. Thus, it may be better to flee, even when one is not absolutely certain that a predator is hiding behind a bush, because the consequences may be particularly harmful. Interestingly, within this framework, the optimal action targets the location where the next fixation will reduce uncertainty the most and not the location that currently looks like the most probable target location. Indeed, both explicit monetary rewards^16^ and implicit behavioral costs^6^ in experimental settings have been shown to influence eye movement choices.

However, Bayesian decision theory is limited to a particular subset of visual tasks, namely tasks that do not involve planning. Repeatedly taking the action with the maximum immediate utility in general may fail in tasks with longer action sequences and delayed rewards depending on the specific task structure. In these cases, an ideal planner based on the more powerful framework of belief MDPs, which contains the ideal observer and the Bayesian decision maker as special cases, is needed to find the optimal strategy. A Markov Decision Process (MDP)^17,18^ is a tuple (*S, A, T, R*, γ), where *S* is a set of states, *A* is a set of actions, *T* = *P*(*s*'|*s*, *a*) contains the probabilities of transitioning from one state to another, *R* represents the reward, and finally, γ denotes the discount factor. In a belief MDP only partial information about the current state *s* is available, therefore a probability distribution over states is kept as a belief state *b*(*s*) = *P*(z*s* | *D*)^19^. The expected reward associated with performing action *a* in a belief state *b*(*s*) is denoted by the action-value function *Q*:

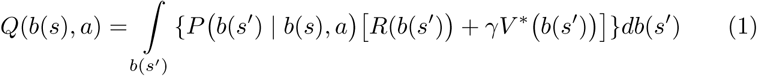
where *V\** (*b*(*s*′)) is the expected future reward gained from the next belief state b(s'). Essentially, what this means is that the value of an action based on the current belief is a combination of the immediate reward and the long term expected reward, weighted by how likely the next belief is under the action. Thus, as the belief about the state of task relevant quantities depends on uncertain observations, actions are influenced both by obtaining rewards and obtaining more evidence about the state of the environment.

In the present study we devised a task that allows probing whether ideal-observer models are sufficient to describe human eye movement strategies. For our visual search task, we derived computational models based on ideal-observer theory as well as on the framework of belief MDPs. Using these models, we specifically created our stimuli such that the two models led either to differ-ent behavioral sequences or to the same behavioral sequences. The rationale for this was to not only show the differences between ideal planer and ideal observer but to also demonstrate that the solutions of both may lead to the same action sequence, depending on the structure of the specific task. Using this experimental paradigm we are able to test whether human eye movement strategies follow the computational principles underlying ideal-observer theory and sequential Bayesian decision making or whether the strategies are planned and future rewards need to be considered (belief MDP).

## Results

**Visual search as planning under uncertainty**. To develop a computational model of visual search as optimal planning under uncertainty it is first necessary to specify the relevant quantities describing the task, i.e. the state representation. In our visual search task (Fig. 1), a suitable candidate for a state representation is the target location and the current location of gaze. However, in general, the exact location of the target is unknown. Therefore, we formalize the probability distribution of the target as a belief state. The action space comprises potential fixation locations and with each action we receive information about the target, update our belief and transition to the next belief state. The reward function is an intuitive mapping between the belief state, which comprises the knowledge about the location of a potential target, and the probability of finding the target.

**Figure 1.**
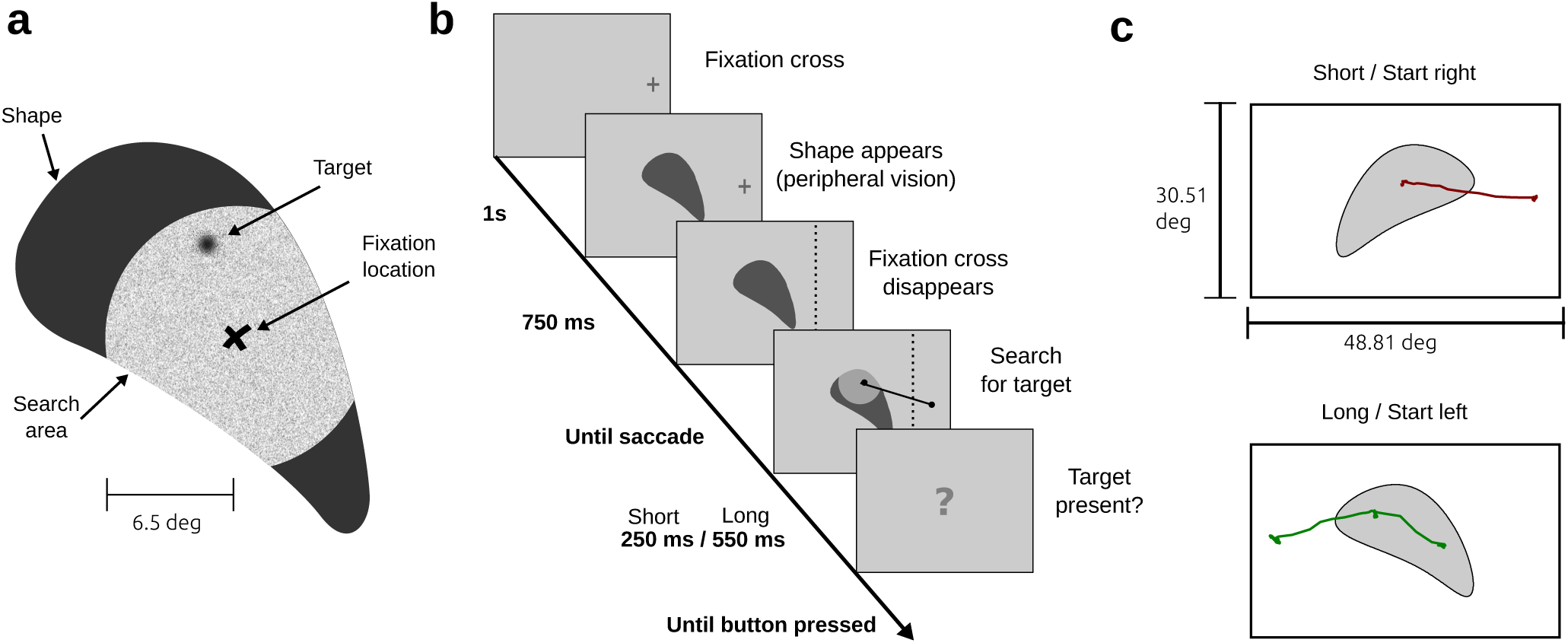
Experimental design. **(a)** Gaze contingent visual search paradigm. Targets were only visible in close proximity to the current fixation location (i.e., inside the search area). **(b)** Procedure for a single trial. Subjects fixated a fixation cross either shown on the left or the right side, respectively. The shape appeared 750 ms prior to the start of the search. The search time was initiated by the participants’ gaze crossing the dotted line. The line, however, was not visible to the subjects. Depending on the condition (short or long) subjects were able to perform one or two fixations inside the shape. **(c)** Raw gaze data is shown for a trial with short search time and initial fixation on the right side (upper panel) and for a trial with long search time and initial fixation on the left side (lower panel). Shapes were mirrored in a counterbalanced design to ensure equal orientation with respect to the initial fixation cross.

How should the actor decide where to look next according to this framework? A policy *π* is a sequence of actions and the optimal policy *π*^*^ comprises actions *a* = arg max*_a_ Q*(*b*(*s*)*,a*) that maximize the expected reward. In tasks comprising sequences of actions, the optimal strategy, the ideal planner, incorporates rewards associated with future actions (*V\**(*b*(*s'*)) into action selection. As a result, the sequence of actions that leads to the maximum total reward is chosen:

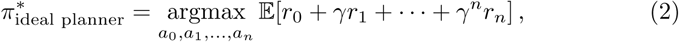
where γ is the discount factor, which controls how much future rewards influence the current action selection.

**Ideal observer as special case of the ideal planner**. If we are only interested in the optimal next action (*γ* = 0) or if there is only a single action to perform equation (1) simplifies to:

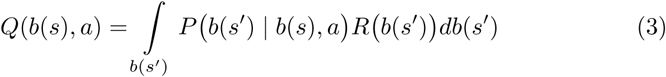
where *P*(*b*(*s*') | *b*(*s*), *a*) is the posterior over relevant quantities in the task and *R*(*s*, *a*) is the cost or reward function. Therefore, if reduced to the next action alone, the ideal planner reduces to the ideal observer with an action selected to maximize task success after the next action. For sequences of actions, the sequential application of the ideal-observer paradigm leads to the action sequence:

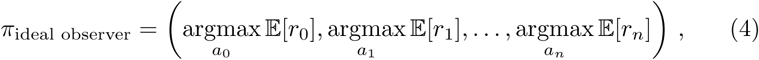
where *a*_0_*,…,a_n_* is the sequence of actions that yields the maximum expected return *r_t_* for each time step t. Whether *π*_ideal_ observer and 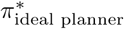 lead to the same action sequence depends on the specific nature of the task. However, in general:

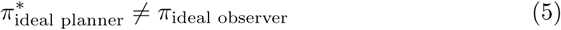
as can be seen in Fig. 2. Ideal-observer approaches only lead to optimal actions if future rewards do not play a role, for example, if only a single action is concerned.

**Figure 2.**
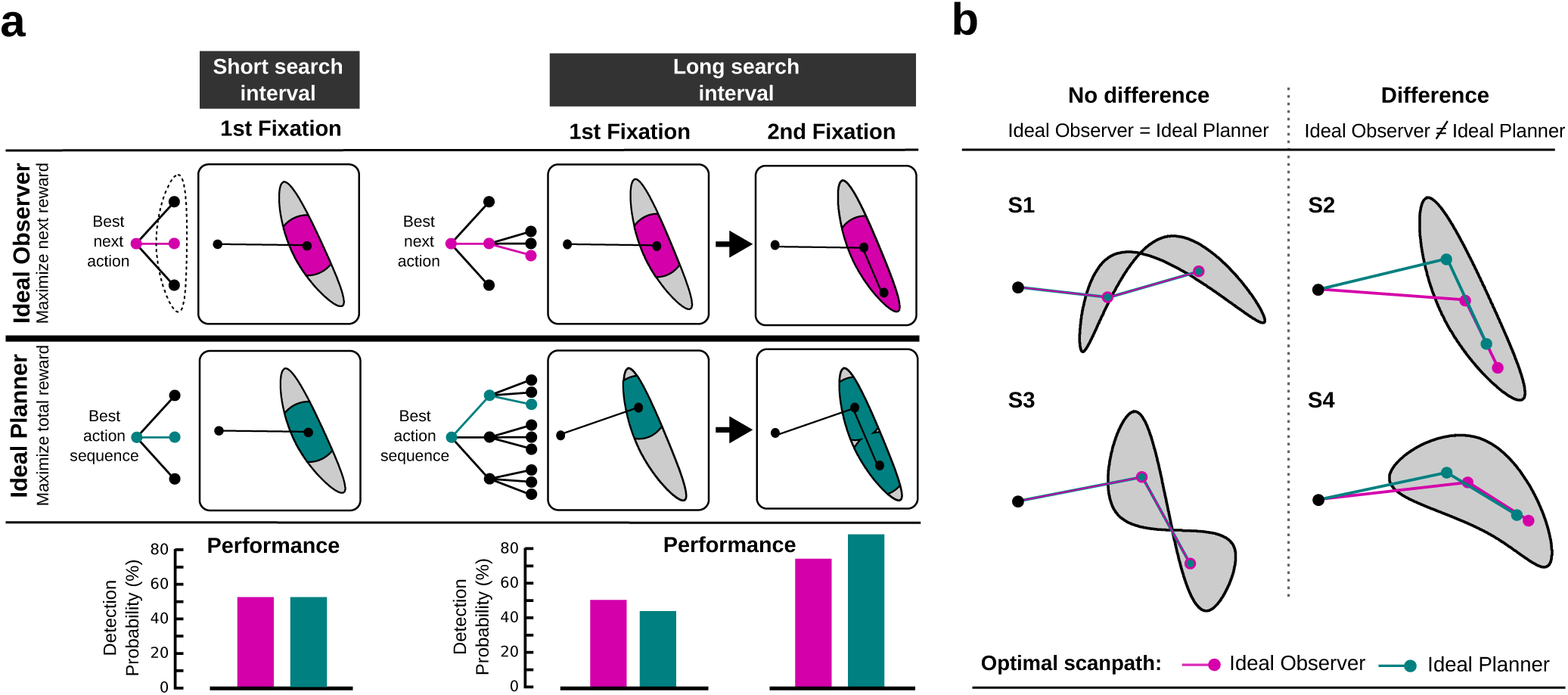
Computational models for visual search. **(a)** Illustration of optimal scanpaths for both models depending on the search time. For the short search interval (left side, one fixation) both models show the same behavior. For the long search interval (right side, two fixations), the ideal observer and the ideal planner differ with respect to the scanpath. While the ideal observer’s next fixation is chosen to maximize the immediate reward (better performance after the first fixation, bottom row), the ideal planner’s scanpath is chosen to maximize performance after two fixations. Computational complexity (depicted as decision trees) is higher for the ideal planner as in the condition with long search intervals all two-fixation sequences are evaluated in order to maximize performance. **(b)** Shapes used in our visual search experiment. For each shape the optimal policy is shown for the ideal observer (pink) and the ideal planner (green). Whether these models lead to different strategies depends on the particular shape. Scanpaths are the same for Shapes S1 and S3, but differ for S2 and S4.

Surprisingly, all of the reviewed computational models for eye movements are myopic, i.e. they choose actions that maximize the immediate reward ^3,20,16,5–7^. In practice, the problem of delayed rewards is circumvented by either investigating only single saccades or by choosing tasks where both policies lead toequivalent solutions. To our knowledge, there exist neither computational models nor empirical data investigating whether humans are capable of planning eye movements. The execution of eye movement sequences has been subject to psychological research and results have shown that the latency of the first saccade was higher for longer sequences of saccades^21^. Also, discrimination performance was enhanced at multiple locations within an instructed sequence of saccades^22^. Further, if an eye movement plan was interrupted by additional information midway the execution of the second saccade was delayed ^23^. Although these results indicate that a scanpath of at least two saccades is internally prepared before execution, no light is shed on whether multiple future fixation locations are jointly chosen to maximize performance in a task.

**Behavioral and model results**. The mean fixation location for each participant separately for all shapes and conditions is shown in Fig. 3a. Also, fixation sequences for the best fit of the ideal observer (right column) and the ideal planner (center column) are depicted. Visual inspection suggests, that the behavioral data is closer resembled by the results of the ideal planner. To test whether eye movements were planned, we compared the first fixation location in the short condition to the first fixation location in the long condition for all shapes. If subjects were capable of performing planning, we expected a difference in the first fixation location for Shape S2 and S4. We used Hotelling’s T-test to compare the bivariate landing positions of the first saccade between the two search intervals (Supplementary Table 1). Indeed, mean target locations for the first saccade were different in Shape S2 and S4. No significant differences, however, were found in shapes S1 and S3. This behavior was well predicted by our ideal planner, but not by the ideal observer. In addition, the direction of the spatial difference of the first fixation location between the search interval conditions followed the course suggested by our ideal planner (Fig. 3b).

**Figure 3.**
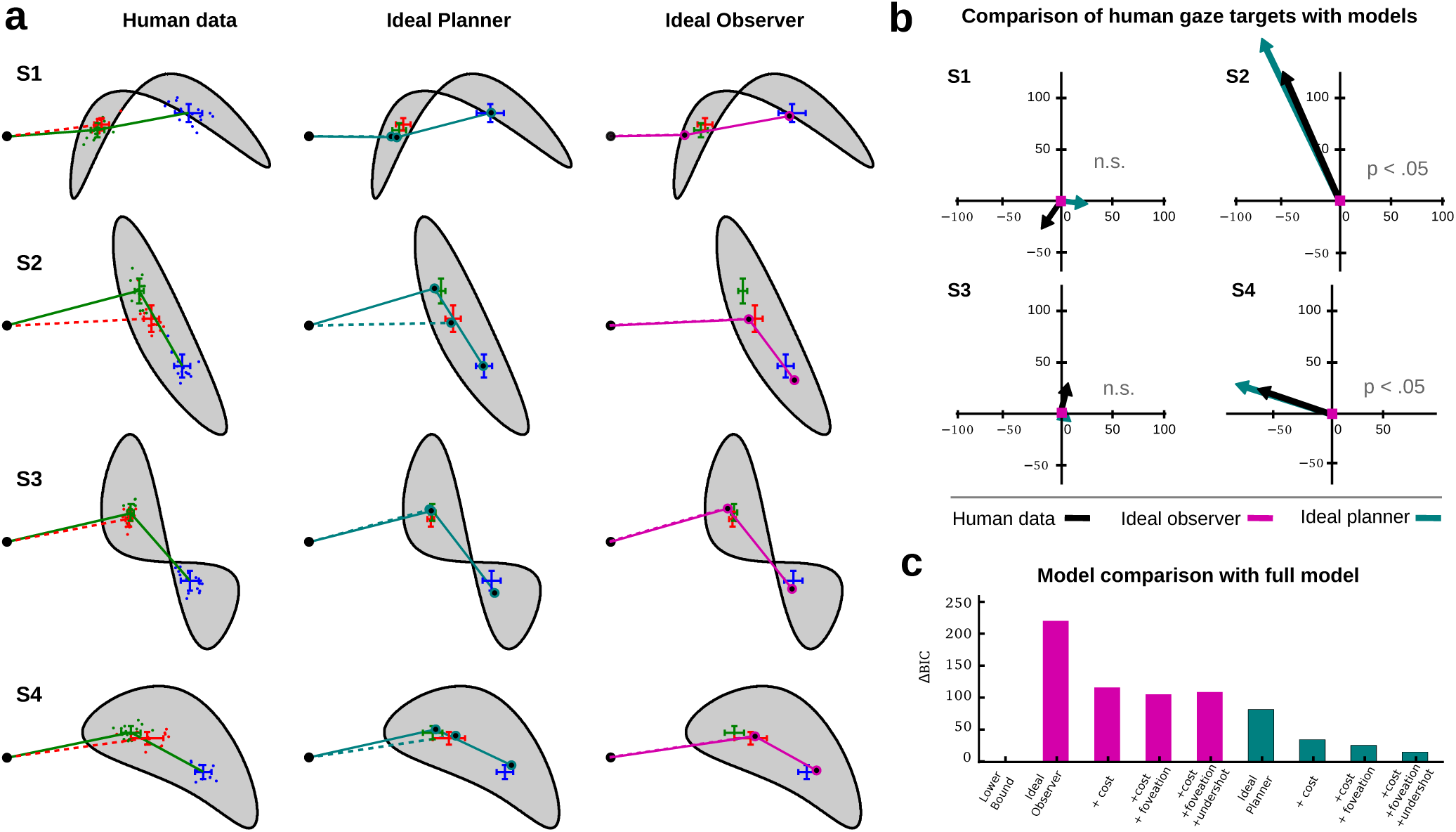
Main behavioral und model results. **(a)** Mean human scanpaths for both conditions (solid lines correspond to long search intervals, dashed lines correspond to short search intervals) are shown in the left column. Colors refer to the condition and the position within the scanpath(red: short search interval, green: first fixation in the long search interval, and blue: second fixation in long search interval). Dots depict mean fixation locations aggregated for each subject individually, error bars show the standard deviation for the fixation location aggregated over all data. The scanpaths suggested by the best fitting models for the ideal planner and the ideal observer are shown in the center and the right column, respectively. Again, solid lines depict the strategy for the long search interval, dashed lines for the short search interval. Global means of the human data are also shown for reference (red, green, and blue). **(b)** Actual and predicted spatial relation of first saccades for all four shapes. Graphs are centered at the fixation location in the short search interval condition. Arrows depict the displacement of the first fixation location in the long search interval relative to the short interval. Arrow color corresponds to the data source. For the ideal observer, the first fixation location is the same for both conditions (indicated by the square centered at (0,0)). **(c)** Difference in BIC between all tested models. The lower bound corresponds to a model directly estimating the mean fixation locations for each shape and condition from the data (3 × 4 means).

**Bounded actor extensions**. We extended both the ideal observer as well as the ideal planner to yield a more realistic model for human visual search behavior, i.e. a bounded actor (see Materials). We added additive costs for longer saccade amplitude (as they lead to longer scanpath duration ^24^ and higher endpoint variability^25^, which humans have been shown to minimize^26^), used foveated versions of the shapes to account for the decline of visual acuity in peripheral vision^27^, and accounted for the often reported fact, that human saccades undershoot their target^28^·^29^. We used the sum of squared errors between our model prediction and our data to compute the BIC for each model. Figure 3c shows the difference in BIC of all models compared to the best model. The lower bound was derived by computing the mean fixation locations directly from the data (3 × 4 parameters). The difference in BIC values between two models is an approximation for the log Bayes factor and a difference Δ*BIC >* 4.6 is considered to be decisive^30^. Results clearly favor the ideal planner over the ideal observer (Δ*BIC* = 138). Crucially, the ideal planner without any parameter fitting still provided a better description of our human data than the ideal observer with all extensions (*ΔBIC* = 27). Further, all model extensions did not only improve our model fit for the ideal planner but were favored by model selection, suggesting that they are needed for describing the eye movement data in our experiment (*ΔBIC* =11 between ideal planner with all extensions and ideal planner without undershot).

Parameter estimates for the saccadic undershot were similar for the ideal observer (4.14 %) and the ideal planner (5.07 %). The influence of the costs for longer saccades was higher for the ideal observer (1.2 DP / Deg) compared to the ideal planner (0.55 DP/Deg). The unit of the costs is detection performance (DP) per degree (Deg) and states, how much performance subjects were willing to give up to shorten saccade amplitudes by one visual degree. We also estimated the radius of the circular gaze contingent search shape centered at the current fixation. Parameter estimation yielded values very close to the true radius and did not improve model quality for neither the ideal planner nor the ideal observer.

## Discussion

It has been unclear whether sequences of human eye movements are planned ahead in time. Prior studies indicate that multiple saccadic targets are jointly prepared as a scanpath and that cueing new targets during execution of eye movements results in longer execution times^21–23^. However, to our knowledge there has been no experimental evidence that eye movements are chosen by considering more than one step ahead into the future. Instead, the ideal-observer paradigm, that models human eye movements as sequential Bayesian decisions has been the predominant approach.

In our study we tested whether the implicit assumptions that accompany the ideal observer are justified. Therefore, we contrasted the ideal observer with the more general ideal planner that was formalized as a Markov Decision Process^18^ with partially observable states^19^. We formalized policies for the ideal observer, only considering the immediate reward for action selection, and for the ideal planner, which also considers future rewards. Next, we derived the specific circumstances under which the models produce different policies. Ultimately, we used these insights to manufacture stimuli that maximized the behavioral differences elicited by the different cognitive strategies and also obtained stimuli that show very similar strategies. Thus, the resulting stimuli were highly suitable for examining which cognitive strategy was adopted by our subjects.

We developed a visual search task where we expected different behavioral sequences depending on the cognitive strategy of our subjects. In particular, we investigated whether subjects adjust their scanpath during visual search dependent on the duration of the search interval. Therefore, we controlled the length of the saccadic sequence. The short search interval allowed subjects to execute a single saccade, while in the long search interval subjects were able to fixate two locations.

Our results suggest that eye movements are indeed planned. Subjects’ scan-path was very well predicted by the ideal planner while showing severe deviations from the scanpath proposed by the ideal observer. Crucially, this was the case even if the sequence required planning. We found fixation locations to be different depending on the duration of the search interval. This difference is only expected under the ideal planner and can not be explain by the ideal observer. Finally, model comparison favored the ideal planner and its extensions over the ideal observer by a large margin. Furthermore, extending our ideal planner model to a bounded planner, we found evidence that subjects traded off task performance and saccade amplitude. Including additive costs for saccades with great amplitude into the ideal planner and accounting for saccadic undershot was best capable of explaining our data further.

Finding and executing near optimal gaze sequences is crucial for many extended sequential every-day tasks^31,32^. The capability of humans to plan behavioral sequences gives further insights into why we can solve so many tasks with ease, which are extremely difficult from a computational perspective. In many visuomotor tasks coordinated action sequences are needed rather than single isolated actions ^33^. This leads to delayed rewards and thus a complex policy is required rather than an action that directly maximizes the performance after the next single gaze switch. Additionally, our findings have implications for future models of human eye movements. While numerous influential past models have not taken planning into consideration^3^·^5^·^6^·^20^, our results indicate that in the case of visual search humans are capable of including future states into the selection of a suitable scan path.

The broader significance of the present results beyond the understanding of eye movements lies in the fact that human behavior in our experiment was best described by a computational model of a bounded probabilistic planning under perceptual uncertainty algorithm. In this framework, sensory measurements and goal directed actions are inseparably intertwined^34,35^. So far, the predominant approach to probabilistic models in perception has been the ideal observer^12,13^, which can be formalized in the Bayesian framework^14,15^ as inferring latent causes in the environment giving rise to sensory observations. Models of eye movements selection have so far used ideal observers^3,5,6^ without planning. Probabilistic, Bayesian formulations of optimality in perceptual tasks^36,37^, cognitive tasks^38,39^, reasoning^40^, motorcontrol^41^, learning^42^, and planning^43^ have lead to a better understanding of human behavior and the quest to unravel, how the brain could implement these computations^44–46^, which are known in general to be intractable^47^. Our results extend the current understanding by demonstrating that planning under perceptual uncertainty is also part of the repertoire of human visual behavior and open up the possibility to understand recent neurophysiological results^48^ within the planning under uncertainty framework.

## Methods

**Participants**. Overall, 16 subjects (6 female) participated in the experiment. The subjects’ age ranged from 18 to 30 years (*M =* 21.8, *SD =* 3.1). Participants either received monetary compensation or course credit for participation. All subjects had normal or corrected to normal vision (four wore contact lenses). One subject stated to have dyschromatopsia, which had no influence on the experiment. Sufficient eye tracking quality was ensured for all data entering the analysis. In each trial a single fixation location (short search interval) or a sequence of two fixation locations (long search interval) entered the analysis. Further, informed consent was obtained from all participants and all experimental procedures were approved by the ethics committee of the Darmstadt University of Technology.

**Task**. In our task subjects searched for a hidden target within irregularly bounded shapes (Fig. 1a). Using a gaze contingent paradigm the hidden target only became visible if a fixation landed close enough 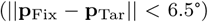. The search area was made explicit by showing the shape’s texture for all points closer than 6.5 to the fixation location. Targets within that area became visible to the participant after a delay of 130 ms. This was done to prevent participants from sliding over the image and instead encourage them to perform distinct fixations. Texture was chosen to reinforce the feeling of looking through the shape (subjects were told to imagine wearing x-ray goggles).

A single trial was as follows (Fig. 1b): Participants fixated a fixation cross that was randomly presented either on the left or the right side of the screen. After 1 *s* the shape was shown in the center of the screen, thus subjects were given access to peripheral information about the shape. Shapes were mirrored if necessary yielding equal distances for left and right starting points. After 750 ms the fixation cross disappeared and participants could initiate the search for the target. Trials in which the first saccade was made while the fixation cross was still visible were dismissed and had to be repeated. It was made transparent to the participants that the search interval started once they made the first eye movement as opposed to when the fixation cross disappeared. After the search interval was over the shape disappeared and participants were asked, whether it contained a target. Overall, shapes contained a target in half the trials. We used two durations as search intervals: a short interval (250 ms) providing enough time for a single saccade and a long interval (550 ms) providing enough time for two saccades. Trials were presented in blocks either containing only short intervals or long intervals, respectively.

**Materials**. Our computational models enabled us to specifically select shapes that facilitate testing our hypothesis. In particular, we identified stimuli that triggered different policies for the ideal-observer model and the ideal-planner model. First, multiple candidates shapes were generated using the following approach: Five points were drawn uniformly in a bounded area (23.24°x 23.24°). Next, a B-spline was fitted to the random points. Finally, the shapes bounded by the splines using the fitted parameters were filled with a texture (white noise). We applied both models to identify shapes that lead to different policies. Overall, four different shapes were used in the experiment (see Fig. 2b). We chose two shapes where optimal behavior requires planning (S2 and S4) and two where it does not (S1 and S3), i.e. where the sequence of eye movements from the ideal observer and the ideal planner coincide. In each category we selected two shapes by visual inspection ensuring that they were similar with respect to the area covered. For display during the experiment the shapes were upscaled with a factor of 1.5 and centered on the monitor such that the center of the shapes bounding box matched the center of the screen.

The target was a circular grating stimulus (0.87°in diameter). Contrast was set in a way that it was easily detected if it was within the visible search radius of the current fixation. The target’s position was generated by randomly choosing a location within the shape.

**Procedure**. After signing a consent form the eye tracker (SMI Red, 250 Hz) was calibrated using a 3 point calibration. Subsequently, subjects completed three to five short training trials (about 1 minute) as part of the experiment instruction. During these training trials it was ensured that the search time was sufficiently long for the individual subject to execute a single saccade in the short condition and two fixations in the long condition, respectively. If necessary the search time was adjusted (between 500ms and 580ms, for the long search interval). Participants were encouraged to ask questions if anything was unclear. After training, participants answered ten questions from a checklist to ensure that they understood the task properly (e.g., when does the search interval start and how many targets can be found at most). Incorrect answers were documented and the correct answers were discussed. After successfully finishing the training, four blocks each containing 100 trials were performed. Thereby, the order of the blocks was either SSLL (two blocks with short search time followed by two blocks with long search time) or LLSS. Participants were randomly assigned to one of the two orders. Eye tracking calibration was renewed before each block.

**Preprocessing**. First, fixations were extracted from the raw gaze signal using the software of the eye tracking device. Overall, 6400 trials (16 participants × 4 blocks × 100 trials per block) entered the preprocessing. 15 trials (0.23 %) were dismissed because the subjects failed to target gaze towards the shape. In these trials, subjects triggered the beginning of the trial by crossing the boundary, however did not engage in visual search. While search time was adjusted to enable subjects to perform a single saccade in the short condition and two saccades in the long condition, respectively, in 17 % of the trials subjects failed to do so. Since we are only interested in comparing the difference between strategies consisting of one or two targeted locations we only used the remaining 5288 trials. Next, we excluded trials where the target was found during executing the search strategy leaving 3145 trials. Clearly, behavior after successfully finding the target is confounded and does no longer provide valid information about the search strategy.

Our analysis and our estimated model parameters rely on mean landing positions aggregated within subjects. Therefore, we need to make sure that the variation in landing positions arises due to saccadic endpoint variability or uncertainties the subject might have about the shape, but not from qualitatively different strategies. Shapes S1 and S3 consist of two separate parts, as a consequence the reward distribution is no longer unimodal across potential gaze targets (see Supplementary Figure 1a). Indeed, qualitatively different strategies in the short condition were found for these stimuli (see Supplementary Figure 1b). Computing mean gaze locations therefore leads to strategies that are qualitatively different from the real data. To further analyze the gaze targets of our participants, we first identified the strategy for each trial using a Gaussian mixture model. We only considered the most frequent strategy (see Supplementary Figure 1c) for both shapes and discarded trials (11.4 %) deviating from the chosen strategy. However, our findings do not depend on the particular choice of strategy as shapes that revealed differences between the ideal observer and the ideal planner (S2 and S4) did not elicit different strategies. The remaining 2785 trials were used for our analysis.

**Model**. Here we derive expressions that implement the general mechanisms of equation (2) and (4) for our visual search task. According to our experimental design participants directed their gaze to suitable locations within a shape in order to decide if a target was present. Depending on the condition, the action sequence in our task comprised one (short condition) or two (long condition) fixation locations. Formally, the greedy policy of the ideal observer (equation (4)) leads to the sequence of fixation locations (*x*_0_, *y*_0_), (*x*_1_, *y*_2_),…, (*x_n_, y_n_*) that maximizes the quality of the decision after each step. In the case of two fixations this leads to:

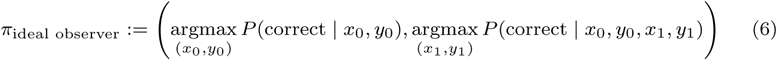
where *x_n_, y_n_* are the coordinates of nth fixation location and *P*(correct|*x_n_,y_n_*) denotes the probability of deciding correctly whether a target is present after the nth fixation.

The non greedy policy of the ideal planner can be derived from equation (2) in a similar fashion. Again, we consider the case of two fixations (LI). Here, the next fixation location is determined by maximizing the reward simultaneously using the next two fixation locations:

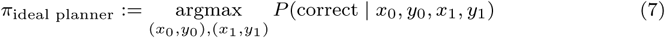

Thereby, (*x*_0_, *y*_0_) is the next location and (*x*_1_, *y*_1_) is the location thereafter. By jointly optimizing the entire sequence of fixation locations the ideal planner is always equal or better compared to the ideal observer. Intuitively, *π*_ideal observer_ and *π*_ideal planner_ yield the same action sequence if the sequence only contains a single action, i.e. a single fixation. Also, the first fixation location of ideal observer is the same for both conditions. Crucially, this is not the case for *π*_ideal planner_. By jointly maximization the reward over the whole action sequence, even the first fixation location can differ between the conditions.

Next, we derive the probability of a correct decision given a sequence of fixation locations since both proposed policies depend on the performance in the task, i.e., the detection probability. The probability of correctly judging the presence of a target is proportional to the area covered by the search. This can be computed as:

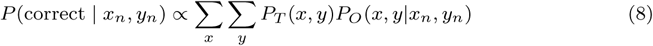
where *P_T_*(*x*, *y*) is the probability that the target is located at (*x*, *y*) and *P_O_* (*x*, *y_n_*|*x_n_,y_n_*) is the probability that the location (*x*, *y*) is covered by the search given that the saccade was targeted at (*x_n_,y_n_*). The former is 1/*N* if (*x*, *y*) lies within the shape and zero otherwise, where *N* is the number of possible target locations. The latter depends on the distance between the saccadic target (*x_n_,y_n_*) and the target location (*x*, *y*). Therefore:

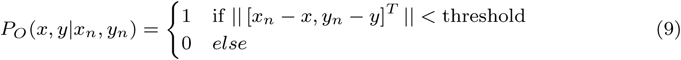
where the threshold is equal to the radius of the search area (6.5°).

**Model extensions**. To take into account known cognitive and biological constraints we need to incorporate several well known characteristics of the human visual system. We introduced costs on the saccade amplitude thus favoring smaller eye movements. As was shown by prior research, greater amplitudes lead to higher endpoint variability^25^ and longer saccade duration^24^. It has further been shown that humans attempt to minimize endpoint variability when execution eye movements ^26^. Therefore, we hypothesized that subjects show a preference for smaller saccade amplitudes. Computationally, we obtain the total reward as a combination of performance and saccade amplitude

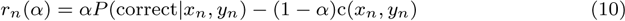
where *c* is a linear cost function returning the amplitude of the saccade. The parameter *α* determines how much detection probability a subject is willing to give up in order to decrease saccade amplitude^6^. It was estimated from the mean fixation locations of our participants using least squares.

Next, the human visual system does not have access to visual content at all locations in the field of view with unlimited precision. We accounted for the decline of visual acuity at peripheral locations. Therefore, foveated versions of the shapes were generated using the known human contrast sensitivity function (see ref. 27, 3, 5, for example). For the first fixation foveation was computed using the initial fixation location of the trial. As it was not computationally tractable to compute foveated images corresponding to the exact location of the first landing position, we approximated it by using the mean fixation location of our subjects instead.

Finally, prior studies have shown that saccades undershot target locations^29^. Initial landing positions are closer to the start location of a saccade. The final target is reached using subsequent corrective saccades. However, in our experiment there is no visible fixation target, therefore corrective saccades might not be present. To account for that we estimated the undershot from our data.

**Code availability**. The codes used for our models are available from the corresponding author on request.

**Data availability**. The data that support the findings of this study are available from the corresponding author on request.

